# Single-Cell Immune Profiling Reveals Non-classical Monocyte-Driven Immunodepression in Tuberculosis Treatment Non-Responders

**DOI:** 10.1101/2025.10.23.684096

**Authors:** Yang Che, Yiqun Xiong, Dongliang Zhang, Yahong Qu, Hao Shen, Weixin Wang, Junshun Gao, ZhenBo Wang, Yi Chen, Zhihong Shen

## Abstract

Sputum culture conversion (SCC) at 2 months is a critical predictor of tuberculosis (TB) treatment outcomes, yet the underlying immune alterations driving treatment non-response remain unclear. We performed single-cell RNA sequencing on peripheral blood mononuclear cells from 8 TB patients, stratified by 2-month SCC status, to characterize cellular heterogeneity and intercellular communication. Treatment non-responders exhibited marked expansion and terminal differentiation of non-classical monocytes with elevated TB progression risk signatures. These cells showed reduced outgoing but enhanced incoming signaling, prominently via IL-16 and TRAIL pathways, potentially amplifying regulatory T cell (Treg)–mediated immunosuppression. CD4⁺ Tregs in non-responders engaged in intensified GALECTIN signaling toward CD8⁺ cytotoxic T cells and mature NK cells, contributing to effector cell exhaustion. NK cell subsets, particularly mature NK c2, displayed increased inhibitory receptor expression and terminal differentiation states. Our findings delineate a dysregulated “non-classical monocyte -Treg -cytotoxic lymphocyte” axis in TB treatment failure, highlighting candidate biomarkers and host-directed therapeutic targets.

## Background

The prevalence of Tuberculosis (TB) and its unfavorable treatment outcome posed a threat to the World Health Organization (WHO) End TB Strategy targets by 2035[1]. Unfavorable treatment outcomes are influenced by multiple factors, including bacteriological characteristics, demographics, clinical history, health care delivery, and socioeconomic conditions [2, 3]. Patients with drug-resistant TB (DR-TB) experience a higher proportion of poor outcomes than those with drug-sensitive TB[4, 5]. Individual risk factors—such as sex, age, education, prior TB treatment, substance use, and resistance to fluoroquinolones—also influence sputum conversion rates (SCR) and the likelihood of achieving sputum culture conversion (SCC)[6, 7].

SCC, defined as two consecutive negative cultures collected at least 30 days apart[8], is a critical early indicator of treatment efficacy and a predictor of long-term cure[9, 10]. Monitoring SCC allows earlier identification of treatment failure, and delayed conversion increases the risk of transmission within communities, highlighting its public health relevance[11, 12]. Despite its clinical importance, the immunological mechanisms that govern SCC remain poorly understood. Host–pathogen interactions are complex and heterogeneous, with significant interpatient variation reflected in radiographic presentation, symptoms, and disease outcomes[13, 14]. It is unclear how these immune interactions and molecular pathways influence the timing or likelihood of sputum conversion.

Single-cell RNA sequencing (scRNA-seq) offers a high-resolution approach to investigate immune heterogeneity and cellular responses at the single-cell level[15]. Previous transcriptional profiling in TB has revealed differences in the state and abundance of immune cell subpopulations early in disease[16], and scRNA-seq has identified functional variation across affected tissues that may act as molecular switches influencing disease progression [17]. However, few prospective studies have systematically linked SCC status with single-cell immune profiling to elucidate the underlying immune alterations.

Considering the crucial role of intercellular communication mediated by cytokines, chemokines, and other signaling molecules in TB pathogenesis and treatment response, we applied scRNA-seq to peripheral blood mononuclear cells (PBMCs) from primary TB patients stratified by 2-month SCC results. This approach aimed to comprehensively characterize the cellular and molecular heterogeneity associated with early treatment response, potentially revealing immunological determinants of SCC.

## Methods

### PBMC Isolation

The scRNA-seq experiment was conducted at the laboratory of Cosmos Wisdom Biotech Co., Ltd. PBMCs were isolated from peripheral blood using LeucoSep tubes prefilled with Ficoll-Paque Plus. Samples were centrifuged at 700 g for 20 min, and PBMCs were collected from the interface, followed by red blood cell lysis and two washes in PBS with 0.04% BSA. Cell suspensions were filtered through 30-µm strainers, and only cells with >90% viability were used for downstream processing.

### Sample Fixation and hybridization

PBMCs were fixed for 24 h at 4 °C and quenched. For each sample, 1 × 10^6 fixed cells were incubated with unique Human WTA Probes for 24 h, washed, counted, and pooled to achieve equal representation. Samples were then processed immediately for GEM generation.

### Single-cell Sequencing

Fixed RNA libraries were prepared using the 10×Genomics Chromium X Instrument and Chromium Fixed RNA Kit. Hybridized samples were partitioned into Gel Beads-in-emulsion (GEMs), barcoded, and ligated probes were pre-amplified. Libraries were quantified with Qubit and sized using Qsep100. Sequencing was performed on an Illumina NovaSeq X Plus (2 × 150 bp).

### Single-cell RNA Statistical Analysis

Raw reads were filtered with fastp, aligned to the human probe set (GRCh38-2020-A) using CellRanger multi v7.1.0, and aggregated across samples. Cells with >200 expressed genes and <20% mitochondrial reads passed quality filtering; mitochondria genes were removed. Doublets were identified and excluded using Scrublet (threshold 0.25)[18]. Genes expressed in fewer than three cells were filtered out.

Normalization, scaling, and regression were performed using Scanpy (v1.9.3). PCA (top 30 PCs) and UMAP were used for dimensionality reduction and visualization. Unsupervised clustering identified cell populations, and marker genes were detected using Wilcoxon rank-sum test (Log2FC >0.25, p<0.05, pct_nz_group>0.1). Clusters of the same cell type were reanalyzed for refined annotation.

### Functional and Pathway Analysis

GSEApy package (version: 1.1.1, https://github.com/zqfang/GSEApy/releases) was used for gene set enrichment analysis[19]. We utilized the scanpy.tl.score_genes function from the Scanpy package to compute feature scores for predefined gene sets [20, 21].To assess the tissue preference of each cell subset, we calculated the ratio of observed to expected cell numbers (Ro/e)[22].

### SCENIC Analysis

To assess transcription factor regulation strength, we applied the Single-cell regulatory network inference and clustering (pySCENIC, v0.9.5) workflow, using the 20-thousand motifs database for RcisTarget and GRNboost[23].

### Trajectory and Differentiation Analysis

Single-cell trajectories were reconstructed using Monocle2 (DDR-Tree)[24], CytoTRACE[25], and Scanpy-based PAGA. Marker genes from Scanpy clustering and normalized expression matrices were used as input. CytoTRACE scores estimated differentiation potential, and PAGA connectivity inferred developmental transitions, validated by lineage-specific marker expression.

### CellChat Cell-Cell Communication Analysis

Intercellular signaling was inferred using CellChat (v1.6.1)[26] based on known ligand-receptor pairs from CellChatDB.human. Outgoing and incoming signaling changes were quantified between experimental groups, and key signaling pathways were identified using the CellChat comparative analysis pipeline.

### Microarray data analysis

Microarray datasets from GEO (GSE40553) were analyzed using CIBERSORT to estimate immune cell subset abundance (perm 100, quantile normalization disabled)[27]. Samples were clustered by Euclidean distance, and gene expression differences between groups were assessed with the Wilcoxon rank-sum test.

## Results

### Single-Cell Transcriptomic Profiles of Immune Cells in Response to Standard TB Treatment

We recruited 8 TB patients, including treatment responders (n=4, Control) and non-responders (n=4, Case), stratified by 2-month SCC status (Fig. 1A). Clinical characteristics including age, sex, treatment history, and drug susceptibility were collected (Supplementary Table 1, Fig. S1A). After quality control, 83,999 PBMCs were retained (40,918 Control; 42,981 Case), with an average of 1,723 genes detected per cell. PCA and UMAP with Leiden clustering identified 17 clusters, which were annotated into six major immune cell types: T cells, NK cells, B cells, myeloid cells, mast cells, and platelets (Fig. 1B, Fig. S1B, Supplementary Table 2). Cell composition analysis indicated higher proportions of monocytes, NK cells, and mast cells in the Case group, although differences were not statistically significant (Fig. 1B–C, Fig. S1C, Supplementary Table 2).

**Figure 1.**
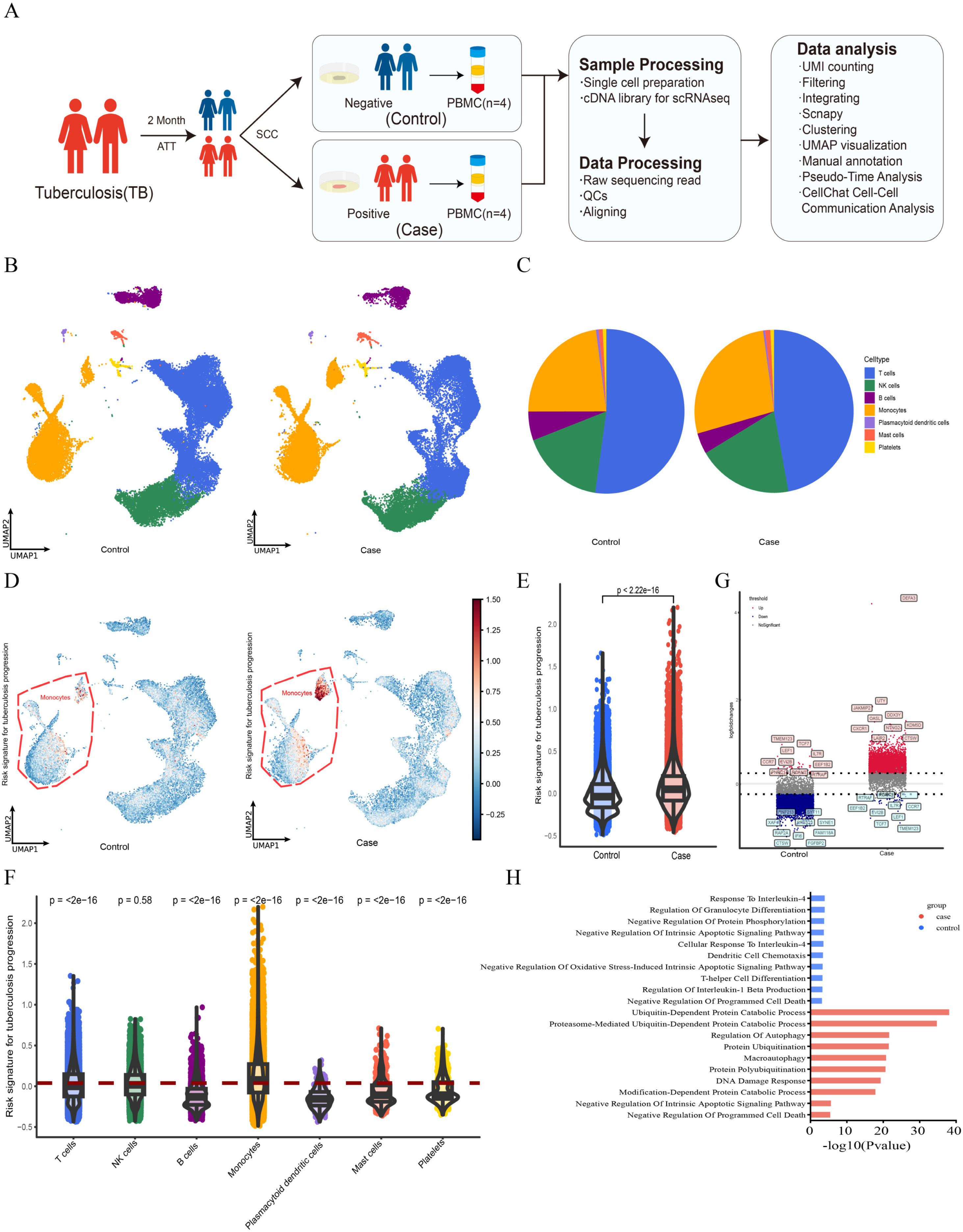
Single-cell transcriptomic landscape of PBMCs from TB treatment responders and non-responders. (A) Schematic overview of the study design. Eight TB patients were recruited and classified into treatment responders (Control, *n* = 4) and non-responders (Case, *n* = 4) based on sputum culture conversion (SCC) status at 2 months after anti-tuberculosis therapy (ATT). (B) UMAP plot of 83,999 PBMCs revealing 17 transcriptional clusters. Cells were annotated into six major immune cell types based on canonical marker genes: T cells, NK cells, B cells, myeloid cells, mast cells, and platelets. (C) Proportions of each major immune cell type in the Control and Case groups. (D) Comparison of TB progression risk scores between Control and Case groups. (E) Violin plots of individual patient risk scores. Statistical significance was assessed using the Wilcoxon rank-sum test. (F) Violin-box-scatter plots showing the distribution of TB risk scores across immune cell types. Each point represents a single cell. The red horizontal line indicates the overall mean risk score across all PBMCs. Statistical comparisons were performed between each cell type and the total PBMC population using the Wilcoxon rank-sum test. (G) Differential gene expression analysis between Control and Case groups across all PBMCs. The top 10 most significantly upregulated genes in each group are labeled. (H) Pathway enrichment analysis showing distinct biological processes activated in the Control and Case groups.

To evaluate disease severity, we applied a previously established TB progression risk signature (Supplementary Table 3)[28]. The Case group exhibited significantly higher risk scores, primarily driven by monocytes (Fig. 1D–F). Expression of GBP2, STAT1, and TAP1 was elevated in monocytes from the Case group (Fig. S1D). Differential gene expression analysis revealed enrichment of T cell–associated genes (TCF7, CCR7) in the Control group, whereas DEFA3 and CXCR1 were upregulated in monocytes, and JAKMP2 and UTY in NK cells in the Case group (Fig. 1G, Supplementary Table 2). Pathway analysis indicated activation of cell survival, anti-apoptosis, protein degradation, and NF-κB signaling in the Case group, while the Control group showed enrichment in immune regulation, cell differentiation, and anti-inflammatory or antioxidant pathways (Fig. 1H). Gene set scoring further confirmed enhanced activation of antigen processing, immune response, and innate and adaptive immune pathways in the Case group, whereas antifungal and antiviral response pathways remained unchanged (Fig. S1E).

### Non-classical Monocytes Are Enriched and Terminally Differentiated in Treatment Non-responders

Myeloid cells exhibited the most prominent transcriptional changes and risk score enrichment in the Case group, prompting subcluster analysis that identified eight subsets (Fig. 2A, Fig. S2A). Among these, non-classical monocytes (NCMs) were relatively enriched in the Case group, with elevated TB progression risk scores (Fig. 2D–E, Fig. S2B–C, Supplementary Table 3). PAGA and Monocle3 trajectory analyses revealed no clear differentiation path, and CytoTRACE indicated that NCMs were terminally differentiated (Fig. 2F, Fig. S2D–E). SCENIC analysis highlighted activation of key transcription factors, including ZNF177, FOXP3, and HES5 (Fig. S2F).

**Figure 2.**
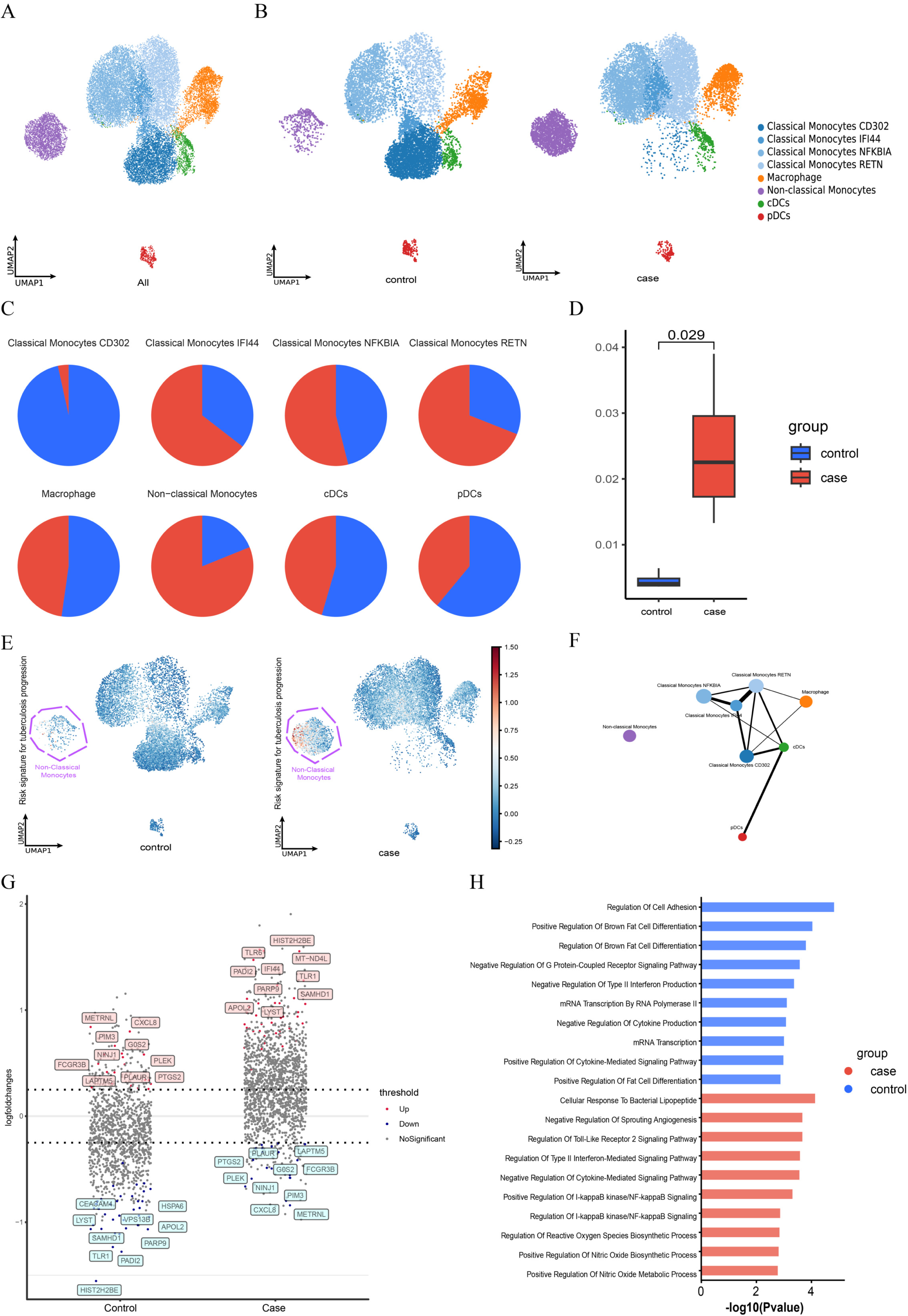
Non-classical monocytes are enriched and terminally differentiated in treatment non-responders. (A) UMAP plot showing the distribution of myeloid cells within the total PBMC population. (B) Proportions of each myeloid subset across individual samples in Control and Case groups. (C) Pie charts comparing the relative proportions of myeloid subsets between Control and Case groups. (D) Box plots showing the relative abundance of non-classical monocytes between Control and Case groups. (E) UMAP plots of TB progression risk scores in monocytes, stratified by group. (F) Trajectory inference analysis of myeloid subsets using PAGA. (G) Differentially expressed genes between non-classical monocytes from Control and Case groups. (H) Gene Ontology (GO) enrichment analysis of biological processes (BP) for differentially expressed genes in non-classical monocytes.

Differential gene expression analysis showed upregulation of TLR6, TLR1, IFI44, and PARP9 in Case NCMs, whereas LAPTM5, CXCL8, and PTGS2 were higher in Control (Fig. 2G). Gene ontology enrichment revealed that Control NCMs were associated with cell adhesion and positive regulation of cytokine-mediated signaling, while Case NCMs were enriched for regulation of Toll-like receptor 2 signaling, type II interferon-mediated signaling, negative regulation of cytokine signaling, and I-κB kinase/NF-κB pathways (Fig. 2H). These findings suggest that NCMs in non-responders are terminally differentiated and exhibit a transcriptional program biased toward innate immune activation and inflammatory signaling.

### Reduced Outgoing and Enhanced Incoming Signaling of Non-classical Monocytes via IL16/TRAIL Pathways

We next examined intercellular communication involving NCMs. Globally, the Case group showed more interactions and stronger overall communication than the Control group (Fig. S3A). However, focusing on NCMs as ligand-sending cells, both the number and strength of outgoing interactions were reduced in the Case group, whereas incoming signals from other cell types were stronger (Fig. S3B–C). Analysis of upregulated interaction pairs revealed significant enrichment of IL16 and TRAIL signaling pathways in Case NCMs, targeting various classical monocyte subsets and CD4⁺ T cells (Fig. 3A–D). These results indicate that NCMs in non-responders receive enhanced regulatory input while exhibiting diminished outgoing signaling, with IL16/TRAIL pathways potentially mediating key cross-talk with adaptive immune cells.

**Figure 3.**
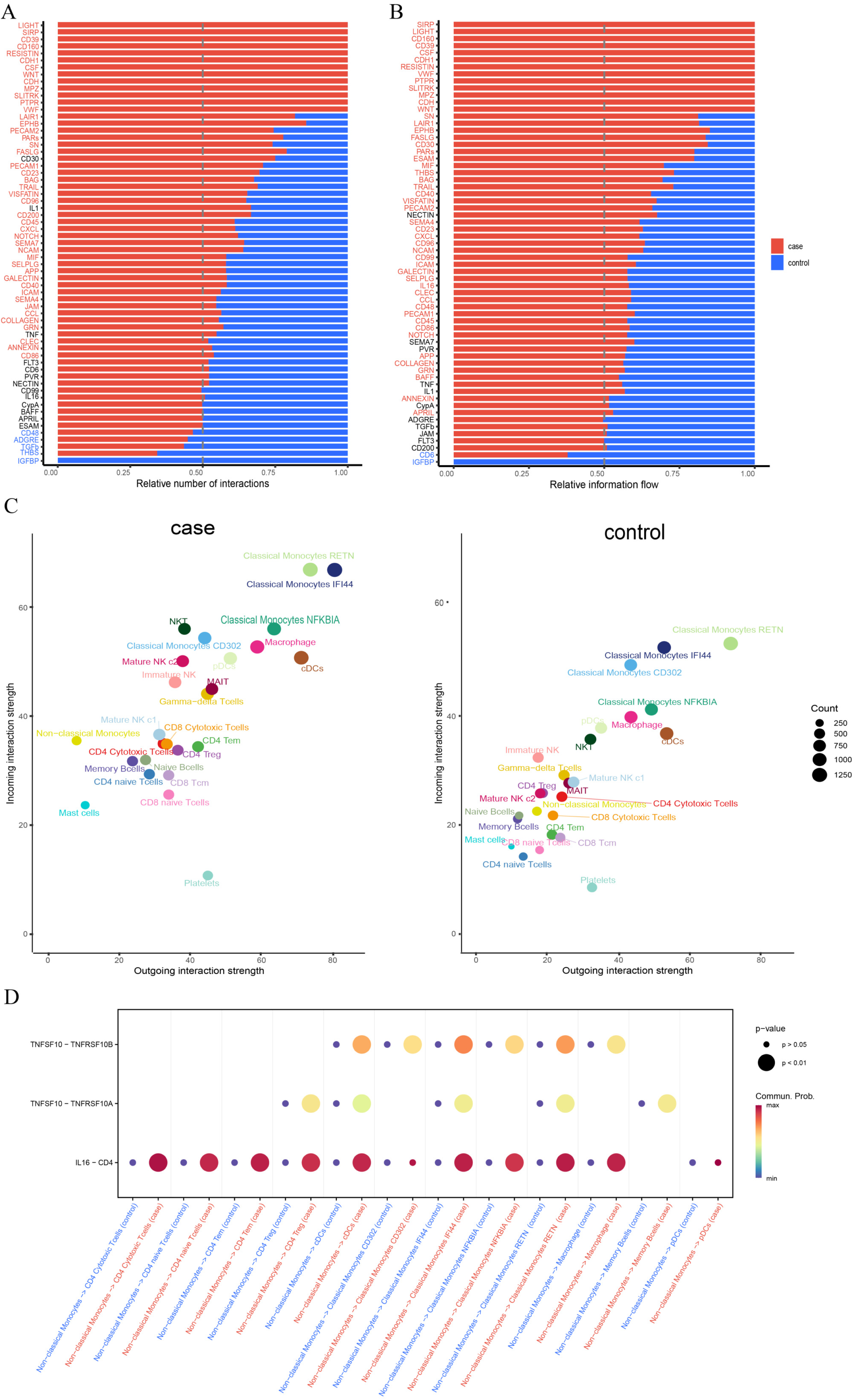
Altered signaling dynamics of non-classical monocytes in Case vs. Control groups. (A) Number of ligand-receptor interaction pairs identified in Case and Control groups. Black indicates no significance; red indicates Case enrichment; blue indicates Control enrichment. (B) Weight of ligand-receptor interaction pairs identified in Case and Control groups. Black indicates no significance; red indicates Case enrichment; blue indicates Control enrichment. (C) Comparison of incoming and outgoing signaling number and strength involving non-classical monocytes as receivers and senders, respectively. (D) Dot plot showing enhanced IL16 and TRAIL signaling pathways with non-classical monocytes acting as ligand sources.

### Cytotoxic T Cell Expansion and LGALS9⁺ Treg-Mediated Immunoregulation in Treatment Non-responders

In addition to altered myeloid signaling, the T cell compartment showed notable changes. Overall T cell abundance was reduced in the Case group, while subclustering revealed a significant increase in CD8⁺ cytotoxic T cells compared to Control (Fig. 4A-D). Functional assessment indicated high cytotoxic potential in both CD8⁺ and CD4⁺ cytotoxic T cells, whereas CD4⁺ regulatory T cells (Tregs) exhibited stronger inhibitory profiles (Fig. 4E, Supplementary Table 3)[29]. LGALS9 expression was significantly upregulated in Case Tregs (log₂FC = 0.85, p_adj = 6.41e-06). CellChat analysis demonstrated that Case Tregs mediated LGALS9-dependent interactions—including LGALS9–P4HB, LGALS9–CD45, and LGALS9– CD44—primarily targeting CD8⁺ cytotoxic T cells, suggesting enhanced Treg-mediated immunoregulation in non-responders (Fig. 4G).

**Figure 4.**
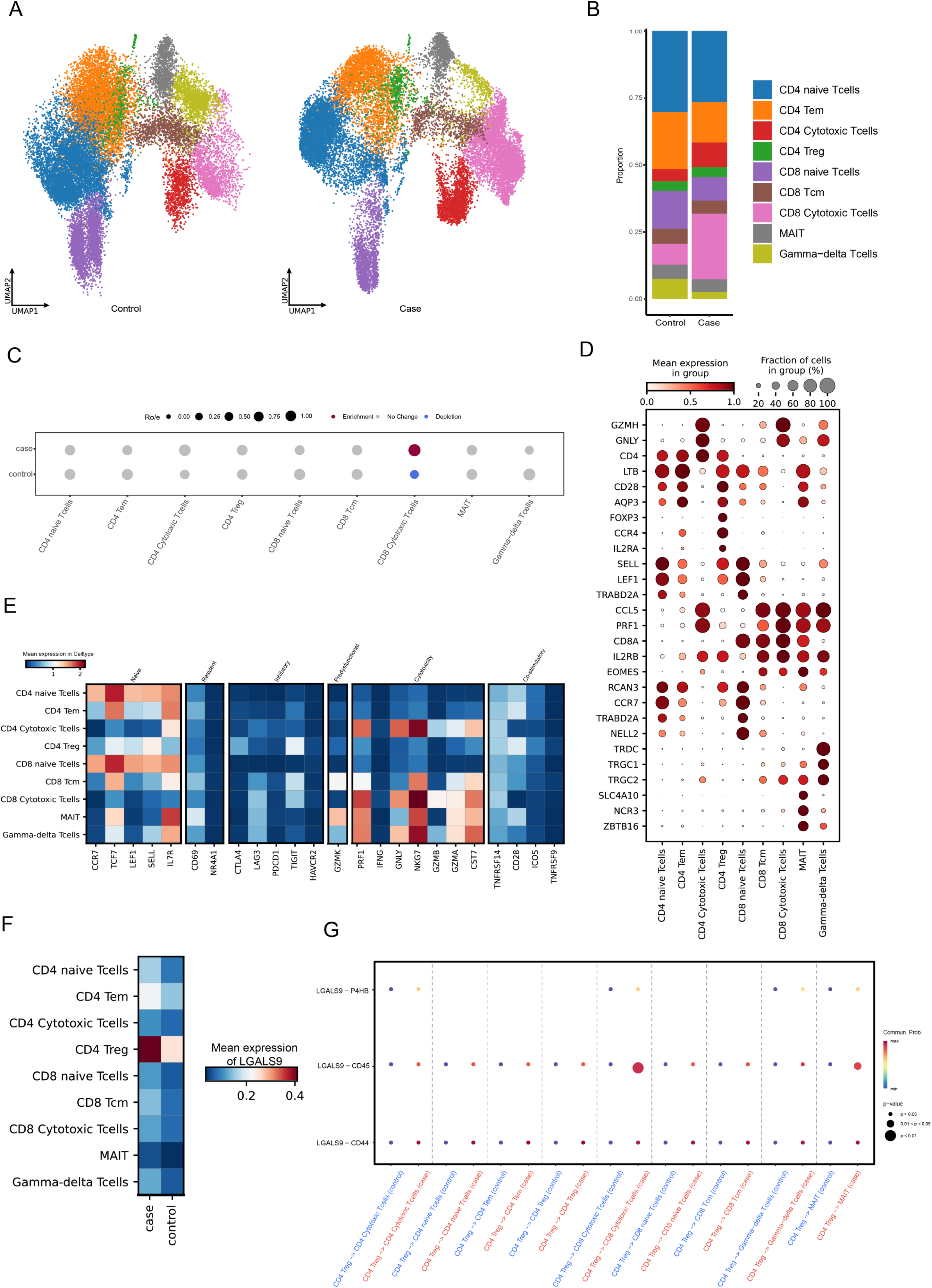
Cytotoxic T cell expansion and GALECTIN signaling-mediated Treg immunoregulation in treatment non-responders. (A, D) Subclustering and annotation of T cell subsets based on canonical markers. (B) Proportion plot showing the distribution of T cell subsets in each group. (C) Dot plot of Ro/e enrichment analysis, with enrichment defined as Ro/e > 1.5 and p < 0.05, depletion as Ro/e < 0.5 and p < 0.05, and no change otherwise. (E) Heatmap showing functional gene expression profiles across T cell subsets. (F) Heatmap displaying LGALS9 gene expression levels in each T cell subset by group. (G) CellChat analysis illustrating GALECTIN signaling from CD4⁺ Tregs (source) to other T cell subsets (receptors).

### Functional Exhaustion and Altered Crosstalk of NK Cell Subsets in Treatment Non-responders

We next examined innate lymphocytes, focusing on NK cells, classified into Immature NK and Mature NK subsets (Fig. 5A, 5D). NKT cells were more abundant in the Case group, and UMAP showed spatial separation of Mature NK c2 between groups (Fig. 5B–C). CellChat analysis revealed stronger GALECTIN signaling, with LGALS9–HAVCR2 interactions between CD4⁺ Tregs and Mature NK c2 significantly enhanced in Case (Fig. 5E). HAVCR2 expression in Mature NK c2 was elevated, while inhibitory receptors such as TIGIT were upregulated in NKT cells. Chemokines CCL5 and CXCR4 were reduced (Fig. 5F–G), indicating functional exhaustion in Mature NK c2 and NKT cells. Pseudotime analysis using Monocle 3 placed these subsets at the terminal end of the developmental trajectory, consistent with a more differentiated or exhausted state (Fig. 5H).

**Figure 5.**
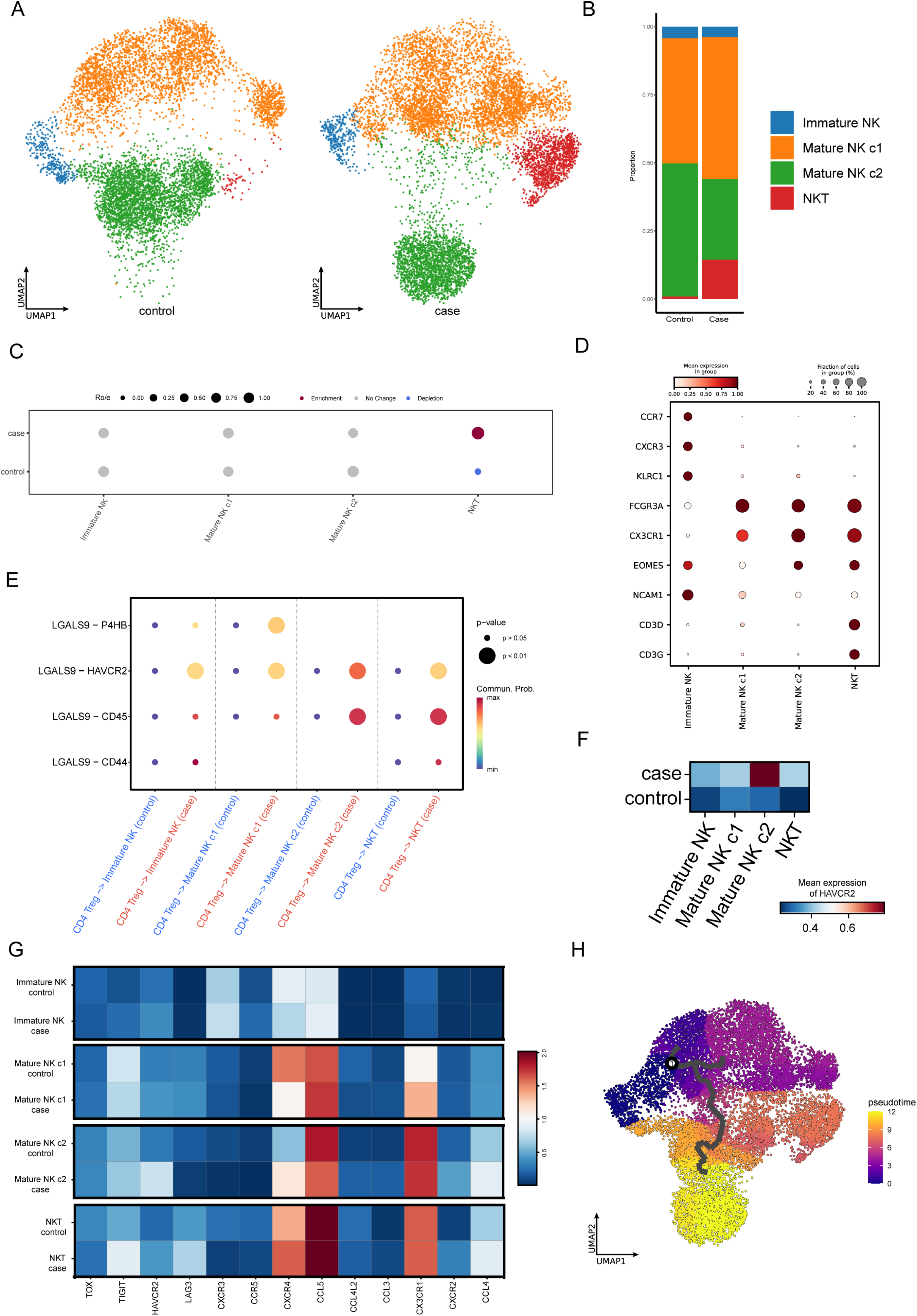
Altered innate lymphocyte composition and GALECTIN signaling in treatment non-responders. (A, D) Subclustering and annotation of NK cell subsets into Immature NK and Mature NK populations based on canonical markers. (B) Proportion plot showing the distribution of NK cell subsets in each group. (C) Dot plot of Ro/e enrichment analysis, with enrichment defined as Ro/e > 1.5 and p < 0.05, depletion as Ro/e < 0.5 and p < 0.05, and no change otherwise. (E) CellChat analysis illustrating GALECTIN signaling from CD4⁺ Tregs (source) to NK cell subsets (receptors). (F) Gene expression levels of HAVCR2 in NK cell subsets across groups. (G) Heatmap showing upregulation of exhaustion markers in NK cell subsets across groups. (H) Monocle 3 pseudotime trajectory analysis visualized by UMAP plot.

### Microarray data validation links non-classical monocyte dynamics to TB treatment response

To validate our single-cell findings, we analyzed GEO dataset GSE40553[27], containing longitudinal transcriptomic data from 35 PTB patients and latently infected individuals (LTB) at baseline, 2 weeks, 2, 4, 6, and 12 months post-treatment. Using CIBERSORT with our single-cell reference, no significant differences in non-classical monocyte proportions were observed between LTB and PTB (p = 0.077) or between baseline and 2 weeks (p = 0.86). However, proportions decreased significantly at 2 months (p = 0.0048) and remained low through 12 months (Fig. 6A–B). Unsupervised clustering of longitudinal trajectories classified patients into three groups. Group 1 maintained higher non-classical monocyte levels at 2–12 months, whereas Groups 2 and 3 declined after baseline. At 12 months, Group 1 showed higher proportions than Groups 2 and 3 (p = 0.052). Gene expression analysis suggested relative enrichment of IL16–CD4 and TNFSF10–TNFRSF10A axes in Group 1, with higher IL16 and trending LGALS9 expression (Fig. S4).

**Figure 6.**
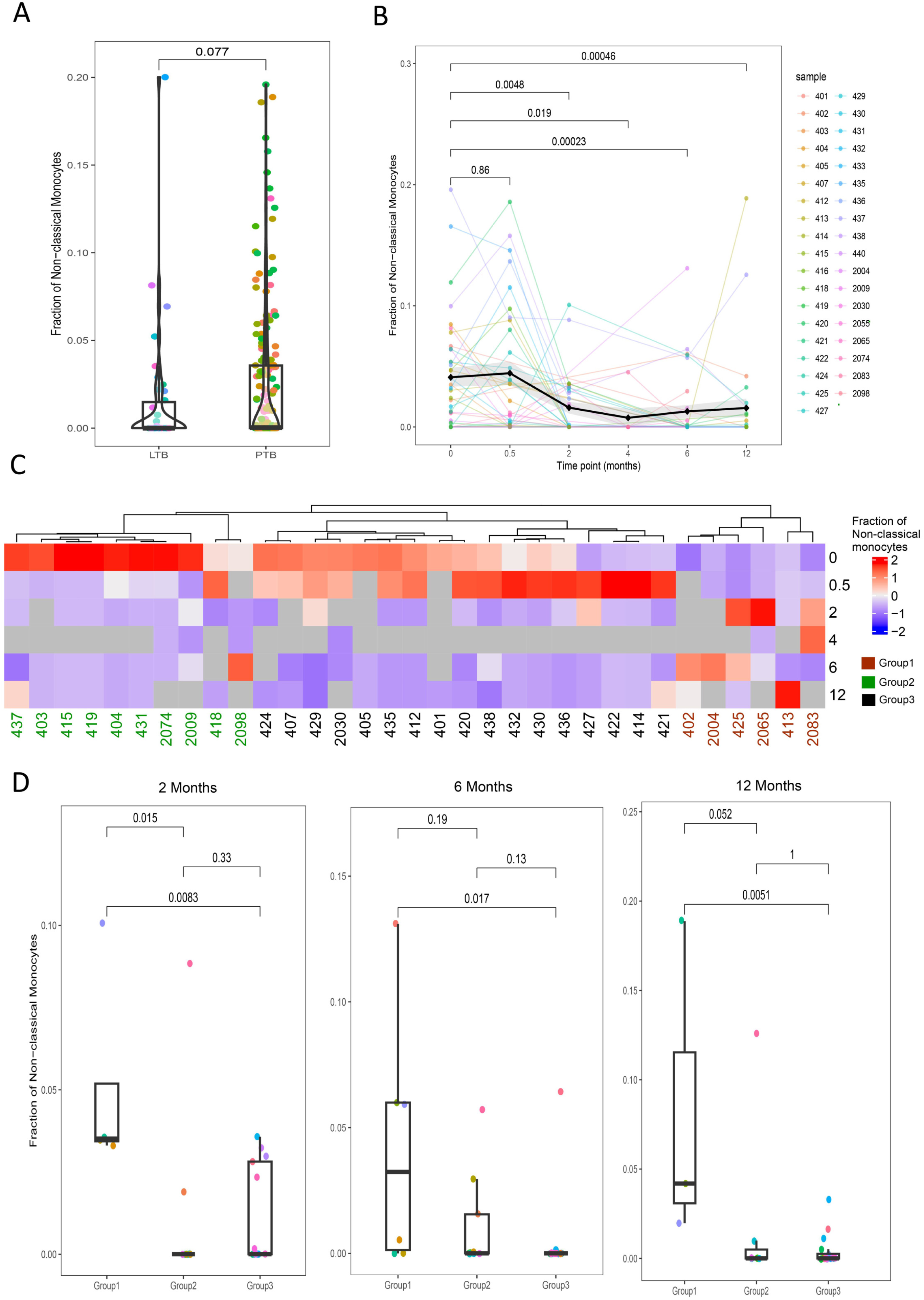
Bulk RNA-seq validation of non-classical monocyte dynamics during anti-TB treatment. (A) Comparison of the proportions of non-classical monocytes between latent tuberculosis (LTB) and pulmonary tuberculosis (PTB) patients, showing no significant difference (p = 0.077). (B) Longitudinal changes in non-classical monocyte proportions during anti-TB therapy in PTB patients. Each line represents an individual patient (colored lines), while the black line denotes the overall mean with the shaded area indicating the standard error. Statistical significance was assessed using the Wilcoxon rank-sum test, revealing no significant change at 2 weeks (p = 0.86), but a significant reduction at 2 months compared with baseline (p = 0.0048), which remained stable at 4, 6, and 12 months. (C) Hierarchical clustering of patients based on standardized non-classical monocyte proportions across all time points. Distinct patient groups were identified, with different colors indicating the clustering results. (D) Comparison of non-classical monocyte proportions at 2, 6, and 12 months across the identified clusters, with statistical testing performed using the Wilcoxon rank-sum test.

## Discussion

Single-cell analysis revealed a striking expansion and phenotypic dysregulation of NCMs in TB patients who failed to achieve sputum culture conversion after 2 months of HREZ therapy (non-responders) (Fig. 1D–F, Fig. 2E, Fig. S2C). Consistent with prior reports linking non-classical monocyte expansion to TB disease severity, we observed marked enrichment of this population in non-responders[30]. The TB progression risk signature score shows that NCMs have been previously associated with advanced disease and impaired bacterial clearance in active TB (Fig. 2C–D, Fig. S2B)[31]. In line with findings by Ong and Hadadi *et al.*, which demonstrated senescence-like features of NCMs driven by NF-κB and IL-1α signaling, our data corroborate these observations (Fig. 1H) [32]. GBP2, STAT1, and TAP1 have previously been proposed as potential biomarkers for MTB infection[33–35]. In our study, we found that the expression levels of these genes in NCMs were higher in treatment non-responders compared with the corresponding responders (Fig. S1D). We show that treatment failure is associated with both quantitative accumulation and distinct regulatory reprogramming of NCMs. Notably, cells from non-responders exhibit a transcriptional signature consistent with terminal differentiation and cellular senescence (Fig. 2G–H, Fig. S2D–E).Intercellular communication analysis showed that NCMs in non-responders receive more incoming signals but emit fewer outgoing signals, acting as signal sinks rather than active communicators (Fig. 3C, Fig. S3B-C). This pattern resembles exhausted or senescent cells that accumulate inhibitory inputs but fail to respond effectively[36]. Similar monocyte dysregulation has been reported in severe TB, where CD14^+^CD16^+^ monocytes correlate with T-cell inhibition, suggesting immune paralysis[37].

A key feature of this dysregulation is the upregulation of IL-16 and TRAIL signaling. IL-16, a pro-inflammatory cytokine and CD4 ligand, is elevated in active TB and promotes MTB survival by inhibiting phagolysosome maturation (Fig. 3D)[38, 39]. NCMs from non-responders express high IL-16, which may recruit Th1 and regulatory T cells but also impair macrophage bacterial killing, fostering chronic inflammation and immune evasion. TRAIL (TNFSF10), also upregulated, can induce apoptosis in infected cells but its dysregulation may cause immune exhaustion and tissue damage by impairing T-cell function and promoting tolerogenic macrophages. Together, the elevated IL-16/TRAIL axis in these monocytes suggests a cytokine imbalance that paradoxically drives inflammation while supporting Mtb persistence[40].

A previous study highlighted the heterogeneity of regulatory T cells between tuberculosis patients and healthy controls, underscoring the critical role of Tregs in immune regulation during TB infection[41]. Our findings further extend this conclusion by demonstrating that in non-responders, CD4+ regulatory T cells exert immunosuppressive effects primarily through GALECTIN signaling, which inhibits the cytotoxic functions of CD8+ T cells and mature NK cell subsets (Fig. 4E-F). Indeed, severe TB is characterized by lymphopenia and widespread immune exhaustion in Th1, CD8+ T and NK compartments[42]. This creates a feedback loop of immune suppression: NCMs fail to activate appropriately and instead secrete suppressive ligands; regulatory T cells expand and inhibit effector cells via GALECTIN; consequently, CD8+ T cells and NK cells become functionally exhausted (Fig. 4G,Fig. 5E,5G). This immune exhaustion severely compromises the host’s ability to clear MTB, thus explaining the persistence of positive sputum cultures despite effective drug therapy.

These mechanistic insights suggest new biomarkers and host-directed targets. Quantifying NCMs expansion or their transcriptomic signature could serve as an early predictor of non-response. For example, a high monocyte or NCMs count at diagnosis has been linked to poor TB treatment outcomes[43]. More specifically, elevated IL-16 or TRAIL levels in blood might herald treatment failure. Importantly, IL-16 and TRAIL are actionable nodes: the recent finding that IL-16 blockade improves TB outcomes in animal models positions it as a candidate Host-Directed Therapy[44].

To further validate the robustness of our findings, we analyzed an independent microarray cohort comprising UK and South African blood transcriptional profiles[27]. In this dataset, NCMs exhibited relatively elevated proportions at baseline (0 months) and after 2 weeks of therapy, but showed a marked decline from 2 months through 12 months of treatment (Fig. 6B). Clustering based on the temporal distribution of non-classical monocyte frequencies identified three distinct patient groups (Fig. 6C). Notably, at 12 months post-treatment, Group 1 displayed significantly different non-classical monocyte frequencies compared with the other two groups (Fig. 6D). Moreover, Group 1 was characterized by consistently higher expression levels of IL16, TNFSF10, TNFRSF10A, and LGALS9 relative to the other groups (Supplementary Fig. S4A–D). We believe that the analysis of this independent cohort provides strong support for our initial observations, reinforcing the notion that non-classical monocyte dynamics and their associated immunoregulatory pathways play a critical role in tuberculosis treatment response.

Together, these findings describe a coherent immunosuppressive network in TB non-responders: dysfunctional NCMs generate IL-16/TRAIL cues that amplify Treg activity, and these Tregs then enforce exhaustion on cytotoxic effector cells. This triangle – “dysfunctional NCMs → hyperactive Tregs → exhausted CD8/NK cells” – can explain the failure of sputum culture conversion (Fig. 7). These immune features not only predict individual treatment failure but also identify patients who are likely to remain infectious for extended periods. Such individuals can act as key drivers of ongoing Mycobacterium tuberculosis transmission, underscoring the urgency for early detection and targeted public health interventions.

**Figure 7.**
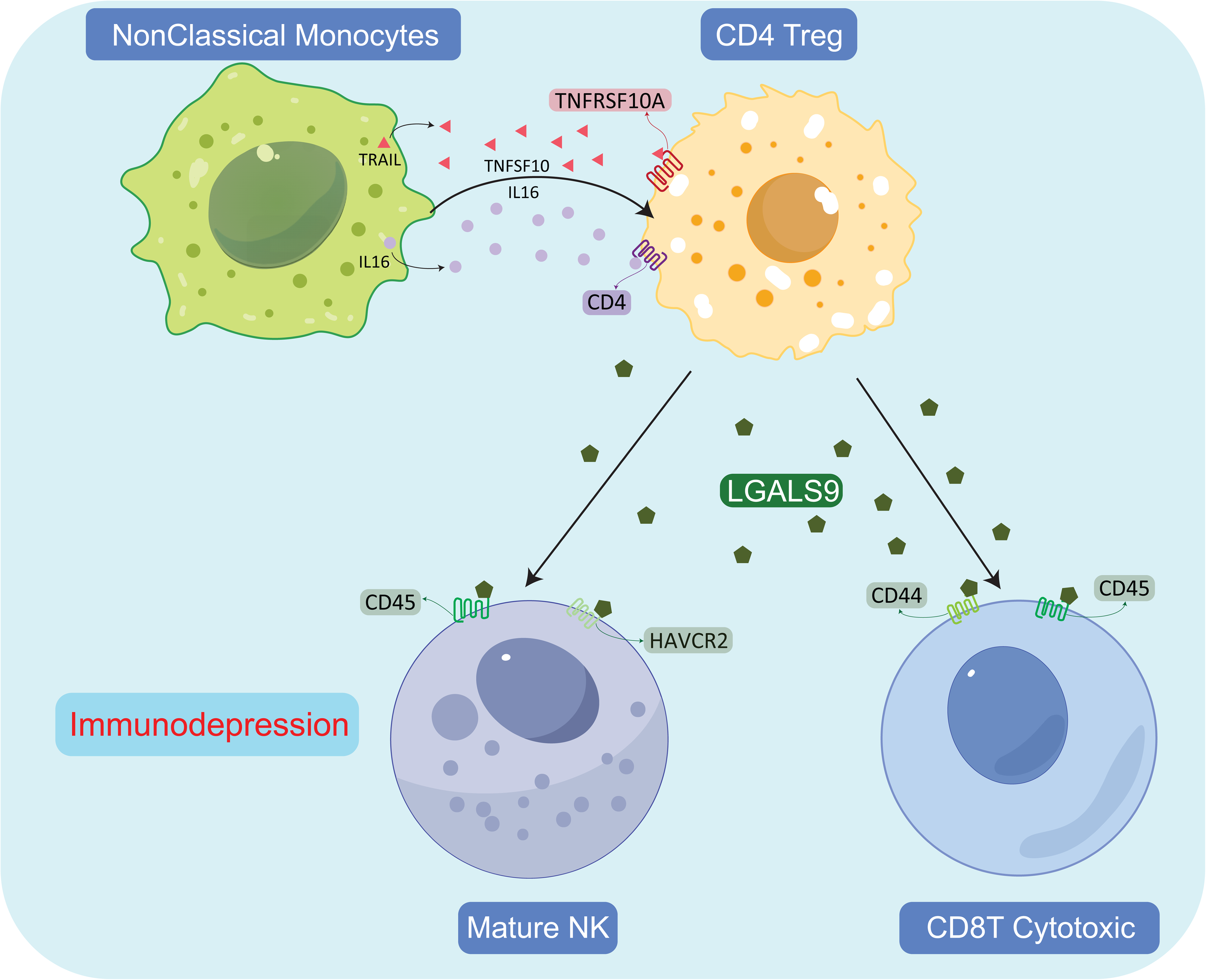
Graphical Abstract. Non-classical monocytes interact with CD4 regulatory T cells (Tregs) primarily through IL16 and TNFSF10 (TRAIL) signaling. Activated Tregs subsequently mediate immunosuppressive effects on mature NK cells and CD8 cytotoxic T cells via LGALS9–HAVCR2 and LGALS9–CD44 axes. These interactions collectively contribute to reduced cytotoxic immune responses in treatment non-responders.

## Declarations

### Ethics approval and consent to participate

This study was approved by the Ethics Committee of the Ningbo Municipal Center for Disease Control and Prevention. All eligible participants who agreed to participate in the program and signed an informed consent form were required to complete a questionnaire and provide specimen for subsequent studies.

### Availability of data and materials

The datasets generated during the current study will be available in the National Genomics Data Center (NGDC) repository (https://ngdc.cncb.ac.cn/) under accession number PRJCA044651 upon completion of submission. Data are currently being submitted and will be accessible following standard NGDC review timelines. Reasonable requests for data during the submission process should be directed to the corresponding author. Bulk RNA-seq raw and normalised microarray data has been deposited with the GEO (GSE40553).

### Authors’ contributions

YaC: Data curation, Writing – original draft.

YX: Formal Analysis, Writing – original draft, Writing – review & editing.

DZ: Formal Analysis, Writing – review & editing.

YQ: Formal Analysis, Writing – review & editing.

HS: Conceptualization, Data curation, Writing – review & editing.

WW: Conceptualization, Data curation, Writing – review & editing.

JG: Conceptualization, Data curation, Writing – review & editing.

ZW: Conceptualization, Writing – original draft, Writing – review & editing.

YiC: Funding acquisition, Supervision, Writing – review & editing.

ZS: Conceptualization, Writing – original draft, Writing – review & editing.

## Acknowledgements

We thank Zhenbo Wang, Hao Shen, Fang Li and Yating Zhang from the Cosmos Wisdom Biotech Co., Ltd. (Hangzhou, China) for providing single-cell related services.

---

**Supplementary Figure S1.**
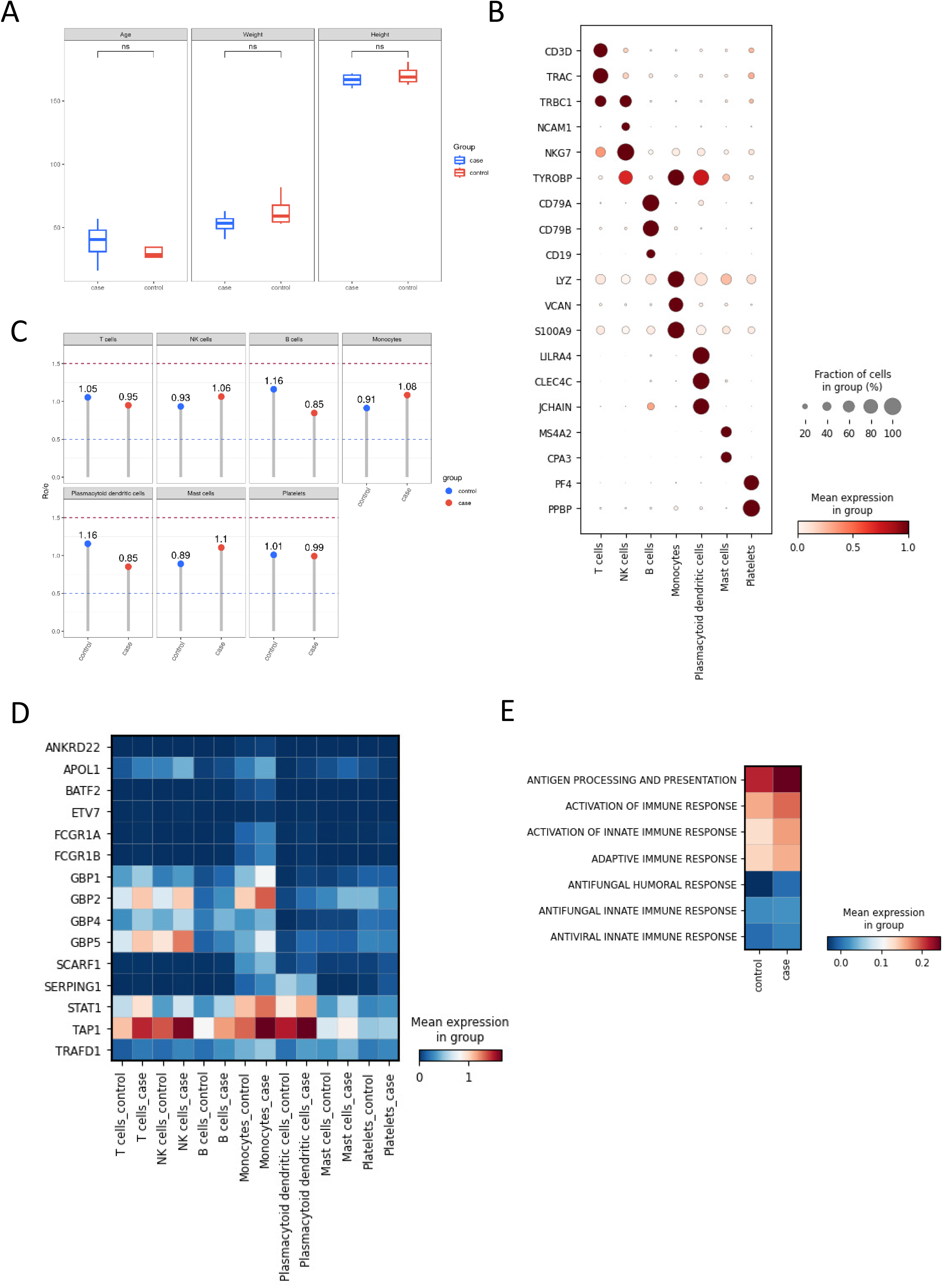
Supplementary analyses supporting the cellular and molecular differences between TB treatment responders and non-responders. (A) Clinical characteristics of patients in Control and Case groups, including demographic and treatment-related variables. (B) Expression of canonical marker genes used to annotate major immune cell types. (C) Relative proportions of immune cell subsets across individual samples. (D) Expression patterns of selected genes in monocytes across groups. (E) Gene set scoring of immune-related pathways comparing Control and Case groups.

**Supplementary Figure S2.**
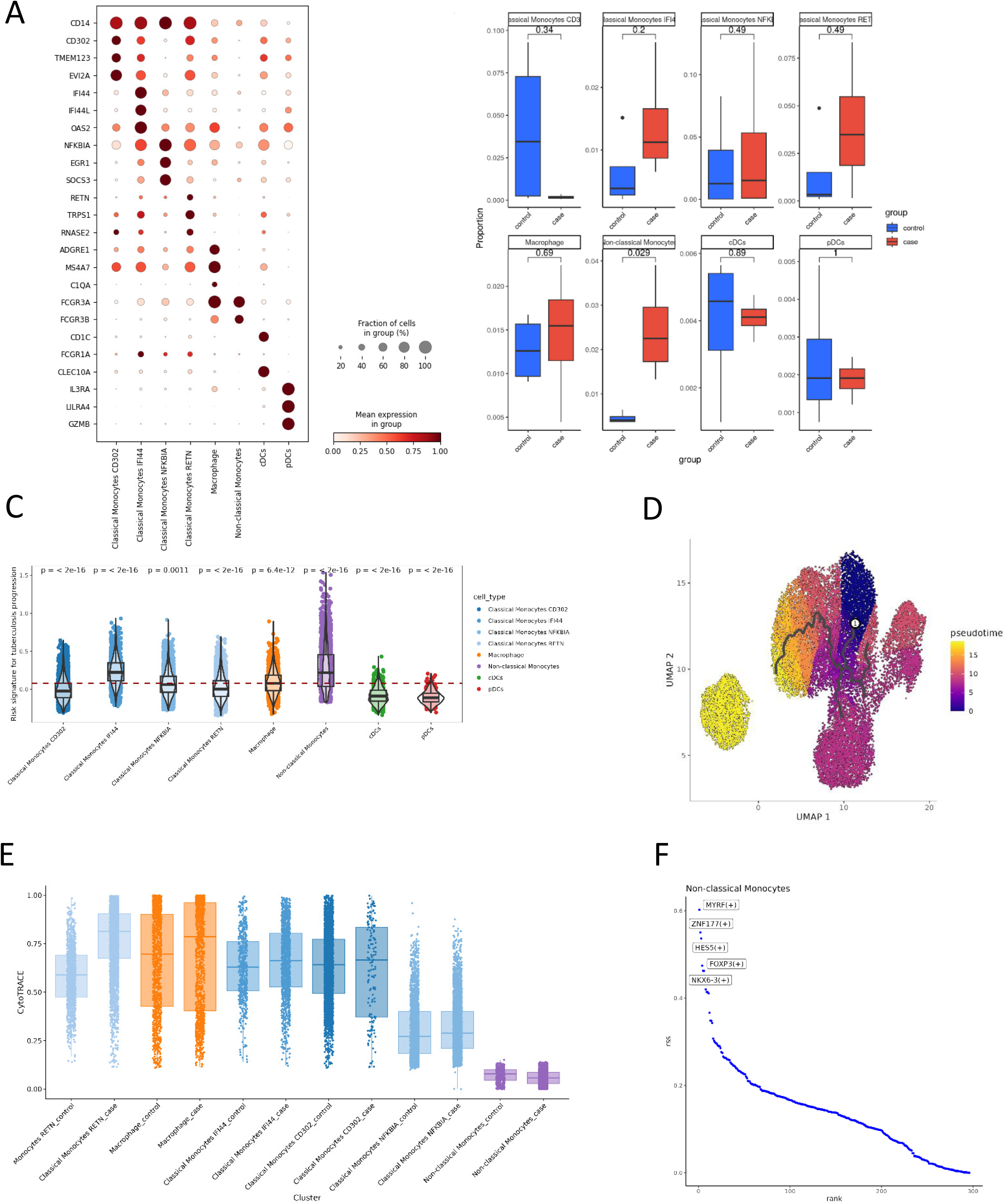
Supplementary analyses of the myeloid compartment and non-classical monocyte differentiation. (A) Dot plot visualization of myeloid subclusters with corresponding annotations. (B) Box plots showing relative frequencies of each myeloid subset across samples. (C) Violin plots showing TB progression risk scores across myeloid subsets in each sample. (D) Monocle3 pseudotime trajectory plots showing differentiation path among myeloid subsets. (E) CytoTRACE analysis indicating differentiation states of myeloid subsets. (F) Dot plot of transcription factor activity inferred by SCENIC analysis in non-classical monocytes.

**Supplementary Figure S3.**
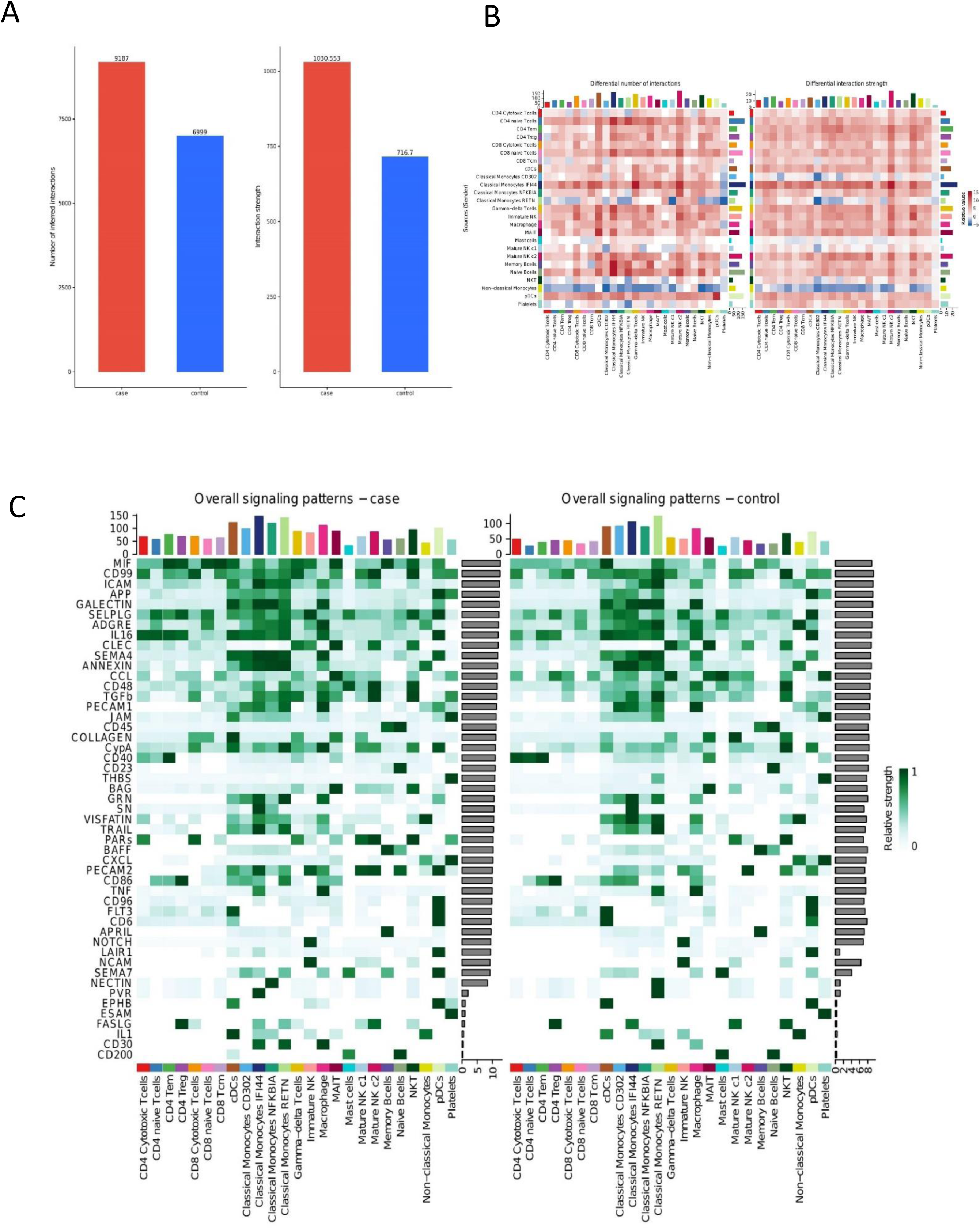
Intercellular communication analysis comparing Case and Control groups. (A) Bar plot showing total interaction counts and overall communication strength across all cell types in Case and Control groups. (B) Heatmap comparing the number and strength of different interactions between Case and Control groups. (C) Heatmap illustrating overall signaling pathway strengths in Case versus Control groups.

**Supplementary Figure S4.**
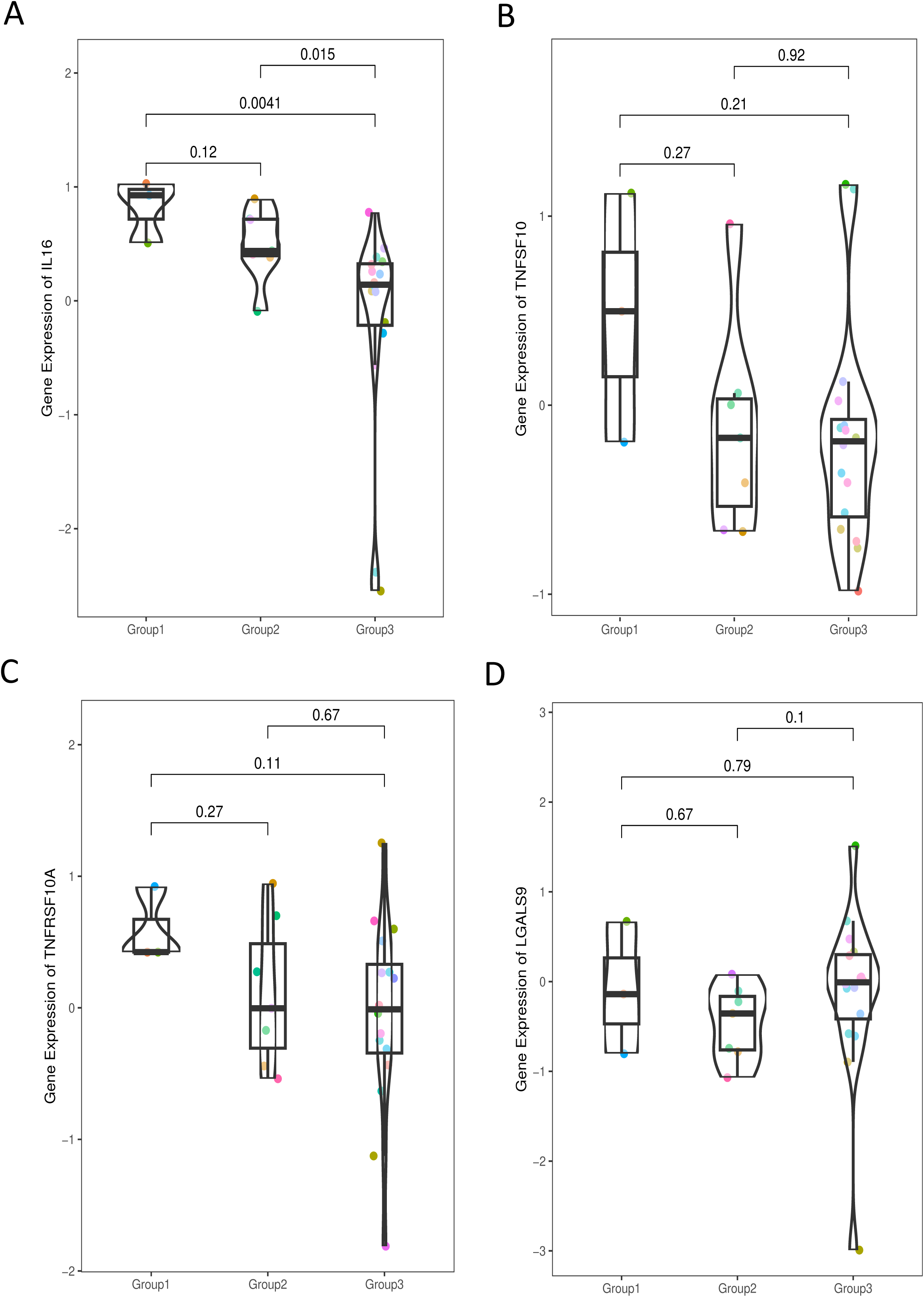
Differential expression of immunoregulatory pathways in non-classical monocytes across patient subgroups. (A–C) Gene expression levels of IL16, TNFSF10, and TNFRSF10A across the three patient subgroups identified by trajectory-based clustering. (D) Expression of LGALS9 across the three subgroups. Statistical comparisons were performed using the Wilcoxon rank-sum test.

**Supplementary Table 1.**

Clinical characteristics of study participants included in the single-cell RNA sequencing analysis. Information includes group allocation (case vs. control), gender, age, anthropometric measurements (weight, height), tuberculosis treatment history, lifestyle factors (smoking, alcohol use), comorbidities, baseline medication regimen, baseline drug resistance profile, baseline microbiology results, 2-month sputum culture conversion status, and overall treatment outcome.

**Supplementary Table 2.**

Single-cell RNA sequencing metadata and differential gene expression results.

Sheet1_MetaData: Per-cell quality control metrics, including total and mitochondrial UMI counts, ribosomal counts, number of detected genes, doublet scores, clustering assignments, and annotated cell type and sub-cell type identities.

Sheet2_CelltypeMarker: Marker genes for each annotated cell type, with associated scores, log fold-changes, p-values, adjusted p-values, and detection rates in group vs. reference.

Sheet3_GroupMarker: Differentially expressed genes between case and control groups, with corresponding scores, log fold-changes, p-values, adjusted p-values, and detection rates.

**Supplementary Table 3.**

Gene sets used in the study for tuberculosis risk signature and immune response analyses. This table lists curated biological pathways and functional gene sets, including Risk signature for tuberculosis progression, Activation of immune response, Activation of innate immune response, Adaptive immune response, Antifungal humoral response, Antifungal innate immune response, Antiviral innate immune response, and Antigen processing and presentation.

